# Delta/theta band EEG differentially tracks low and high frequency speech-derived envelopes

**DOI:** 10.1101/2020.07.26.221838

**Authors:** Felix Bröhl, Christoph Kayser

## Abstract

The representation of speech in the brain is often examined by measuring the alignment of rhythmic brain activity to the speech envelope. To conveniently quantify this alignment (termed ‘speech tracking’) many studies consider the overall speech envelope, which combines acoustic fluctuations across the spectral range. Using EEG recordings, we show that using this overall envelope can provide a distorted picture on speech encoding. We systematically investigated the encoding of spectrally-limited speech-derived envelopes presented by individual and multiple noise carriers in the human brain. Tracking in the 1 to 6 Hz EEG bands differentially reflected low (0.2 – 0.83 kHz) and high (2.66 – 8 kHz) frequency speech-derived envelopes. This was independent of the specific carrier frequency but sensitive to attentional manipulations, and reflects the context-dependent emphasis of information from distinct spectral ranges of the speech envelope in low frequency brain activity. As low and high frequency speech envelopes relate to distinct phonemic features, our results suggest that functionally distinct processes contribute to speech tracking in the same EEG bands, and are easily confounded when considering the overall speech envelope.

**Highlights:**

- Delta/theta band EEG tracks band-limited speech-derived envelopes similar to real speech
- Low and high frequency speech-derived envelopes are represented differentially
- High-frequency derived envelopes are more susceptible to attentional and contextual manipulations
- Delta band tracking shifts towards low frequency derived envelopes with more acoustic detail

## 1. Introduction

A central notion governing the neurobiology of speech has been that rhythmic brain activity aligns to the regularities of speech, particularly in the delta (1 – 4 Hz) and theta (4 – 8 Hz) bands. This alignment has been termed “speech entrainment” or “speech tracking” (Obleser and Kayser, 2019) and both, serves as a powerful tool to study speech encoding, but also promotes a fundamental role of rhythmic brain activity for speech comprehension and production (Giraud and Poeppel, 2012; Obleser and Kayser, 2019; Poeppel and Assaneo, 2020). Experimental signatures of speech tracking in M/EEG or ECoG recordings are often obtained by relating the neural signal to a simplified signature of the acoustic energy: the latter is often defined as the overall or broadband speech envelope and is computed by averaging individual band-limited envelopes along the spectral dimension (Ahissar et al., 2001; Aiken and Picton, 2008; Doelling et al., 2014; Kayser et al., 2015; Keitel et al., 2018; Meyer, 2018; Nourski et al., 2009). However, relevant information is not distributed homogeneously along this spectral dimension: lower frequencies often carry more information about the majority of phonemes and are more relevant for comprehension (Elliott and Theunissen, 2009; Erb et al., 2020; Monson et al., 2014). Also, the auditory system can selectively track spectrally-distinct acoustic streams (Brodbeck et al., 2019; Mesgarani and Chang, 2012; O’Sullivan et al., 2019) and many auditory cortices segregate sounds along a tonotopic organization. If the experimentally observed speech tracking signatures arise from such processes, combining individual envelopes along the spectral dimension would effectively blend distinct neural representations into a single experimental measure. Hence, we conjecture that the use of the broadband envelope to study the encoding of speech and other natural sounds may obscure possibly distinct underlying neural processes and might provide an oversimplified picture.

To investigate this hypothesis, we studied the representation of synthetic stimuli derived from natural German speech in the human EEG. The stimuli were created by applying amplitude-envelopes derived from limited bands of the spectral dimension of natural speech to band-limited noise carriers and hence captured essential aspects of the envelope dynamics of natural speech. Quantifying the alignment of rhythmic components of the EEG to the acoustic envelope (Giordano et al., 2017; Kayser et al., 2015; Park et al., 2016) we then compared the cerebral tracking of such individual band-limited speech envelopes across a number of experimental conditions designed to address a number of questions, as explained below. Across experimental conditions we manipulated the spectral match between individual envelope and carrier bands, the number of simultaneously presented envelope-carrier pairs, and participants’ attention to specific envelope-carrier pairs. In particular, by varying the number of simultaneously presented envelope-carrier pairs, the stimulus material varied along a continuum from individual envelopes to multi-band noise-vocoded speech.

With this approach we were able to investigate the following questions: First, we asked whether envelopes extracted from lower and higher spectral bands (c.f. Fig. 1) are tracked similarly in the EEG, i.e. over the same electrodes and in the same EEG bands (Fig. 2). Second, we asked whether this tracking is sensitive to the match between carrier frequency and the spectral range covered by the envelope, motivated by relevance of the natural match between carrier frequency and envelope band for comprehension as shown by natural and spectrally rotated speech (Molinaro and Lizarazu, 2018; Peelle et al., 2013) (Fig. 3). Third, we probed whether and for which envelope bands tracking is affected by the presence of additional acoustical envelope information (Fig. 4) and by attention (Fig. 5), similar as the known attentional effects for the tracking of natural speech (Cusack et al., 2004; Kerlin et al., 2010; Lakatos et al., 2008; Rimmele et al., 2015; Teoh and Lalor, 2020). Fourth, we investigated the tracking of individual spectrally-limited envelopes and the superposition of multiple envelope-carrier pairs to probe which tracking signatures covary with the signal’s complexity and participant’s comprehension (Fig. 6). And last but not least, we investigated the statistical regularities of acoustic landmarks that supposedly give rise to speech tracking in order to illustrate the necessity of studying the tracking of spectrally limited envelopes (Fig. 7).

**Fig. 1.**
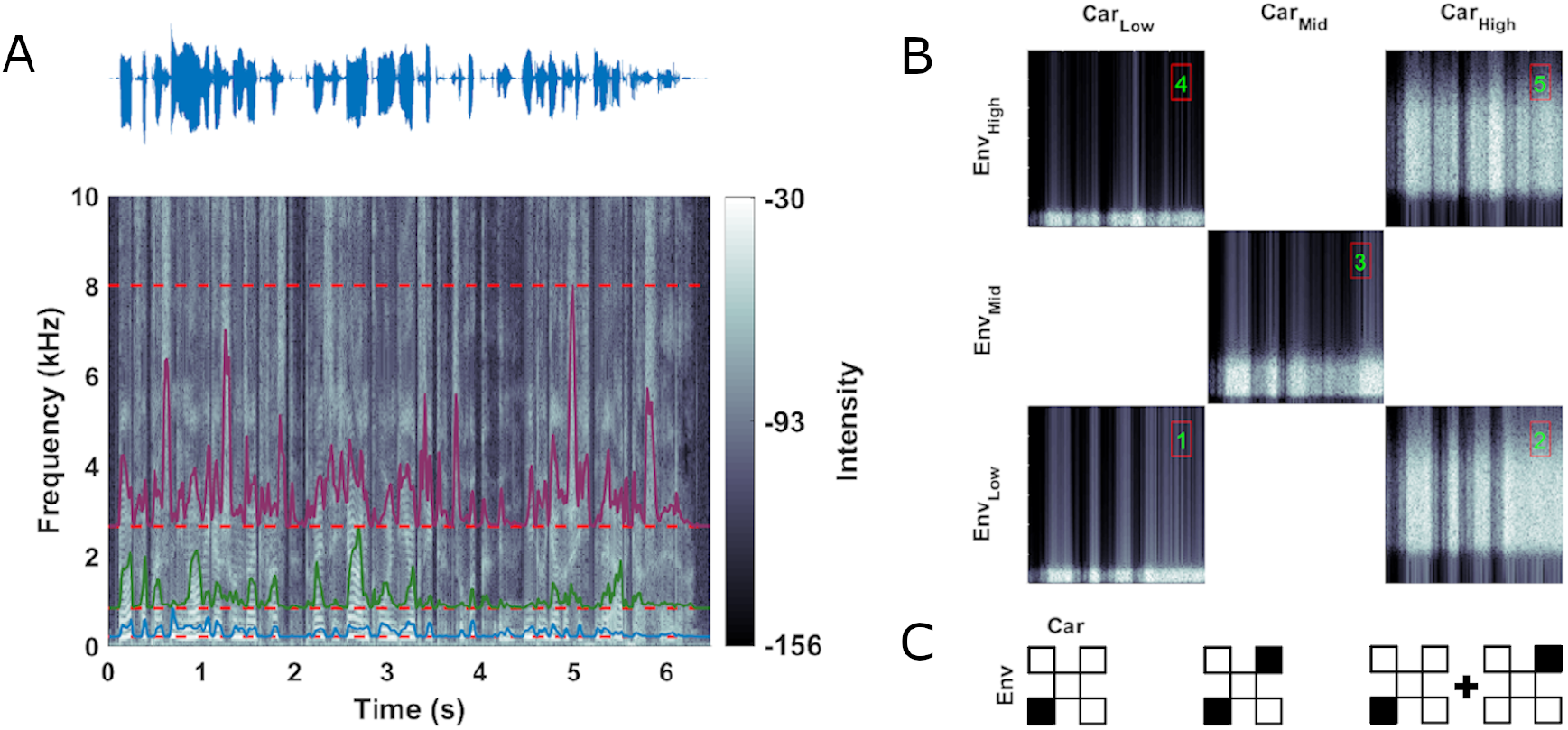
Stimulus design. (**A**) Acoustic waveform (top) and spectrogram (bottom) of one example sentence used to create the stimulus material for the present study. Colored outlines show amplitude envelopes extracted from (here) three logarithmically spaced frequency ranges between 0.2 and 8 kHz (red dashed lines). (**B**) Stimuli were envelope-modulated band-limited noises differing in the frequency range used to create the noise and the frequency range from which the speech envelope was derived and used to modulate the noise. The figure shows 1 second snippets from stimuli used in Block 1. The spectral ranges covered by the carrier noise and the spectral range from which the envelope was extracted were paired across conditions as shown by the matrix (low: 0.2 – 0.83 kHz, mid: 0.83 – 2.66 kHz and high: 2.66 – 8 kHz). Conditions 1, 3 and 5 (green numbers) reflect natural pairings where the frequency band of the carrier was modulated with the envelope derived from the same band, while conditions 2 and 4 reflect unnatural pairings. (**C**) Icons are used in the following figures to illustrate how distinct envelope and carrier bands were used to create a specific stimulus condition in the experiment, or to define a factor for data analysis. Envelope-carrier pairs presented in the experiment are marked as filled boxes in the checkerboard. Depending on the experimental blocks, this could be individual (left) or simultaneously presented pairs (middle). For some data analysis we averaged experimental conditions as indicated with a plus symbol between icons (right), or contrasted these as indicated by a minus sign.

**Fig. 2.**
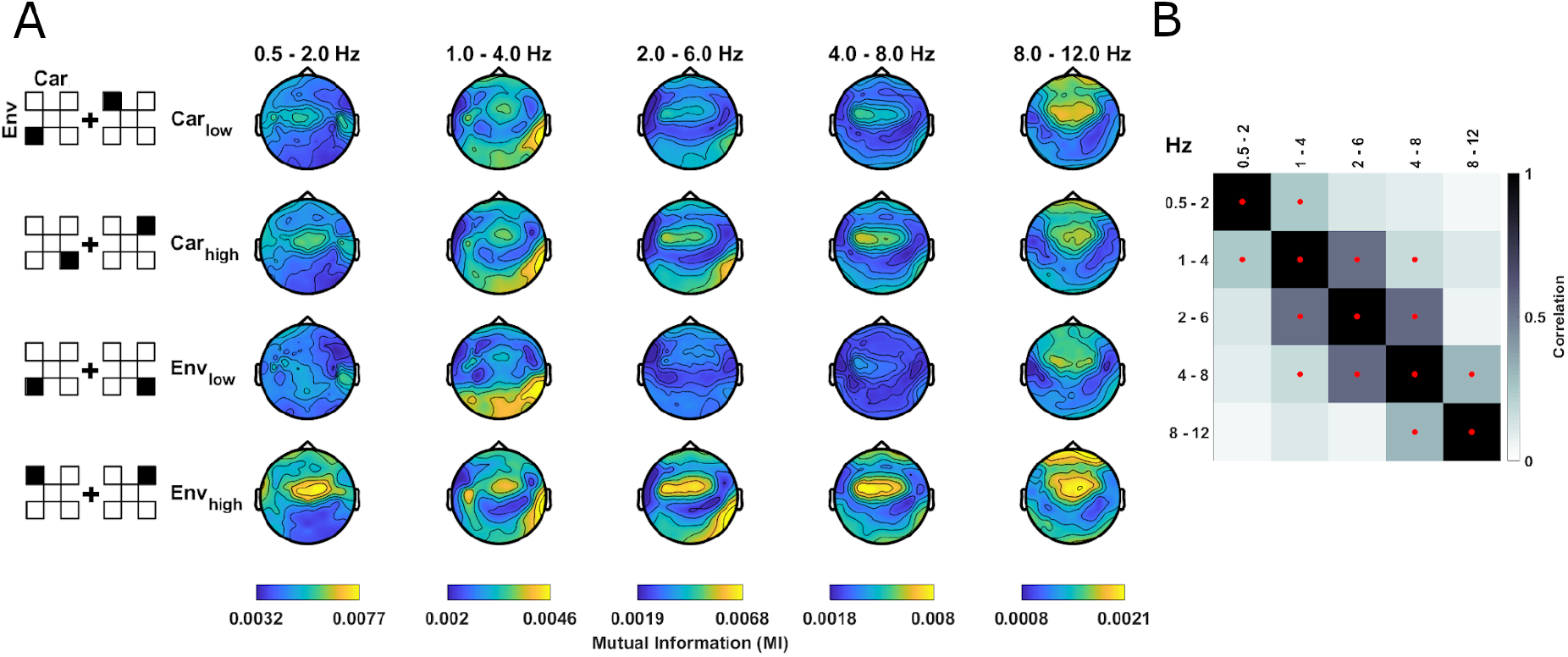
Specificity and spatial topographies of envelope tracking. (**A**) Condition-averaged topographies of envelope tracking, quantified as mutual information (MI), separately for the factors low/high carrier bands (Car_low_, Car_high_) and low/high envelope ranges (Env_low_: 0.2 – 0.83 kHz, Env_high_: 2.66 – 8 kHz) in block 1. To create the analysis factors, the MI values were averaged across the respective other dimension (e.g. over envelope bands for Car_low_, Car_high_), as indicated by the pictograms on the left. These tracking signatures were significant for each envelope and carrier when compared with a bootstrap distribution reflecting the null hypothesis of no systematic relation between EEG and stimulus (see main text). Color ranges are fixed within each EEG band. (**B**) Spatial similarity (Pearson correlation) of group-level MI topographies between EEG bands (averaged over conditions). Red dots indicate a significant (p<0.01) one-sided group-level bootstrap test against zero.

**Fig. 3.**
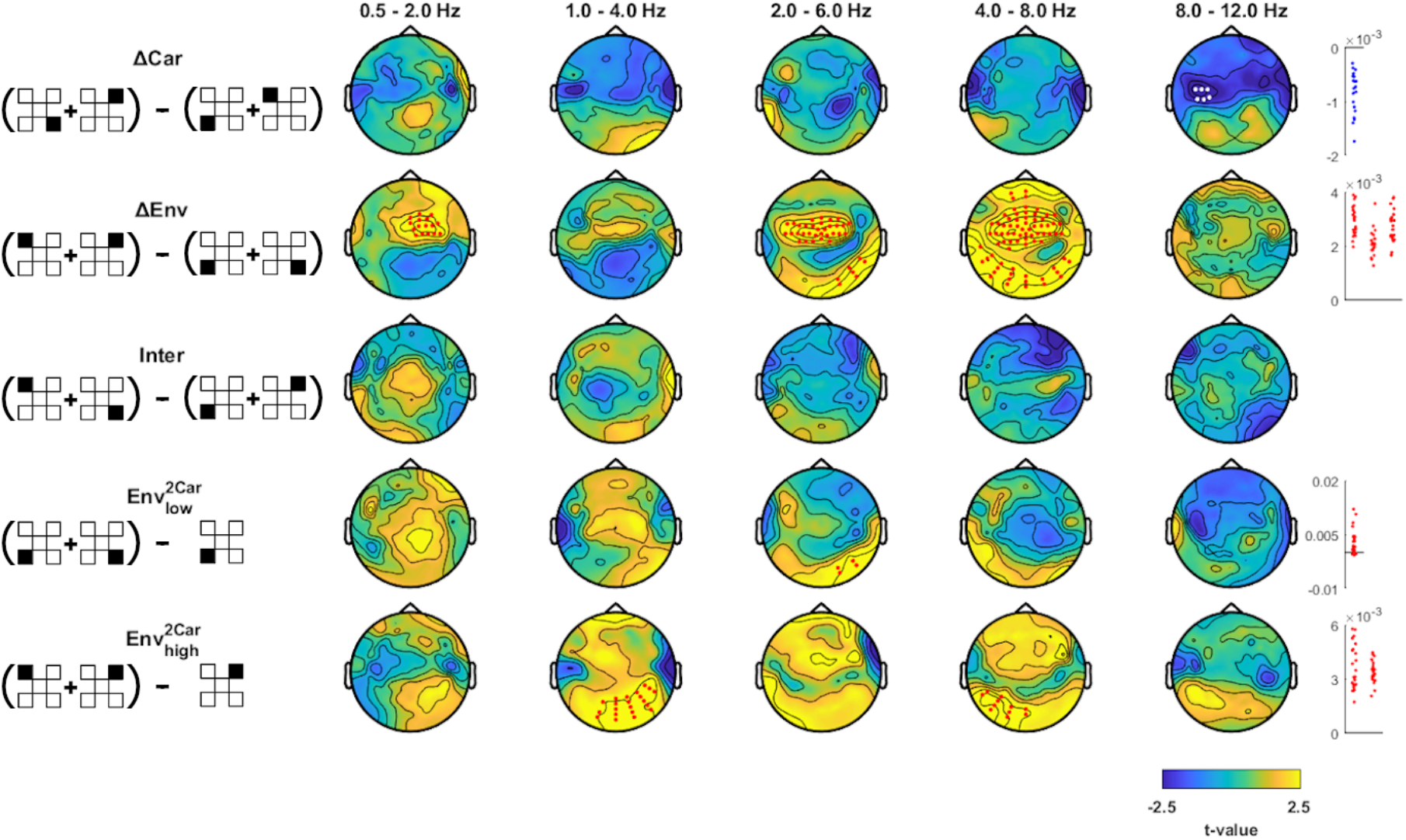
Effects of carrier frequency and envelope band on envelope tracking. Differences in MI values between factors Car_high_ minus Car_low_ (ΔCar), Env_high_ minus Env_low_ (ΔEnv), unnatural minus natural pairings (Inter), and the enhancement of envelope tracking when presented with two minus one carrier (Env_low_^2Car^, Env_high_^2Car^). Pictograms on the left indicate which experimental conditions (black filled squares) were combined into analysis factors and contrasted (c.f. Fig. 1C). Topographies show the electrode-wise t-values for the group-level differences between factors of interest. Red and white dots mark positive and negative clusters (derived using cluster-based permutation statistics corrected across electrodes and EEG bands, p<0.01). Panels on the right show differences in MI values between factors for individual participants within each cluster (red for positive and blue for negative clusters).

**Fig. 4.**
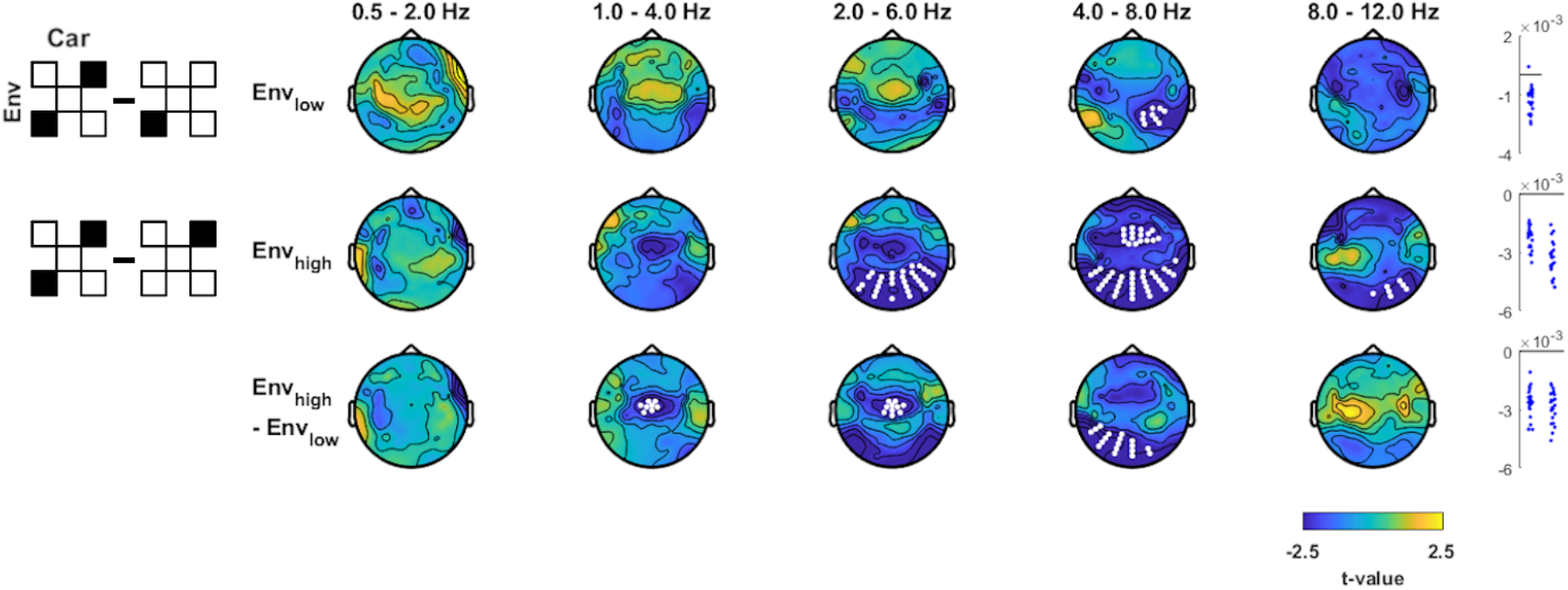
Effects of contextual information. Tracking of one individual envelope-noise pair when a second envelope-noise pair is presented (using a distinct carrier) minus tracking of just the individual envelope. Pictograms on the left indicate which conditions (black filled squares) were contrasted. Topographies show the electrode-wise t-values for the group-level statistics. Red and white dots mark positive and negative clusters (derived using cluster-based permutation statistics, p<0.01). Panels on the right show differences in MI values between factors for individual participants.

**Fig. 5.**
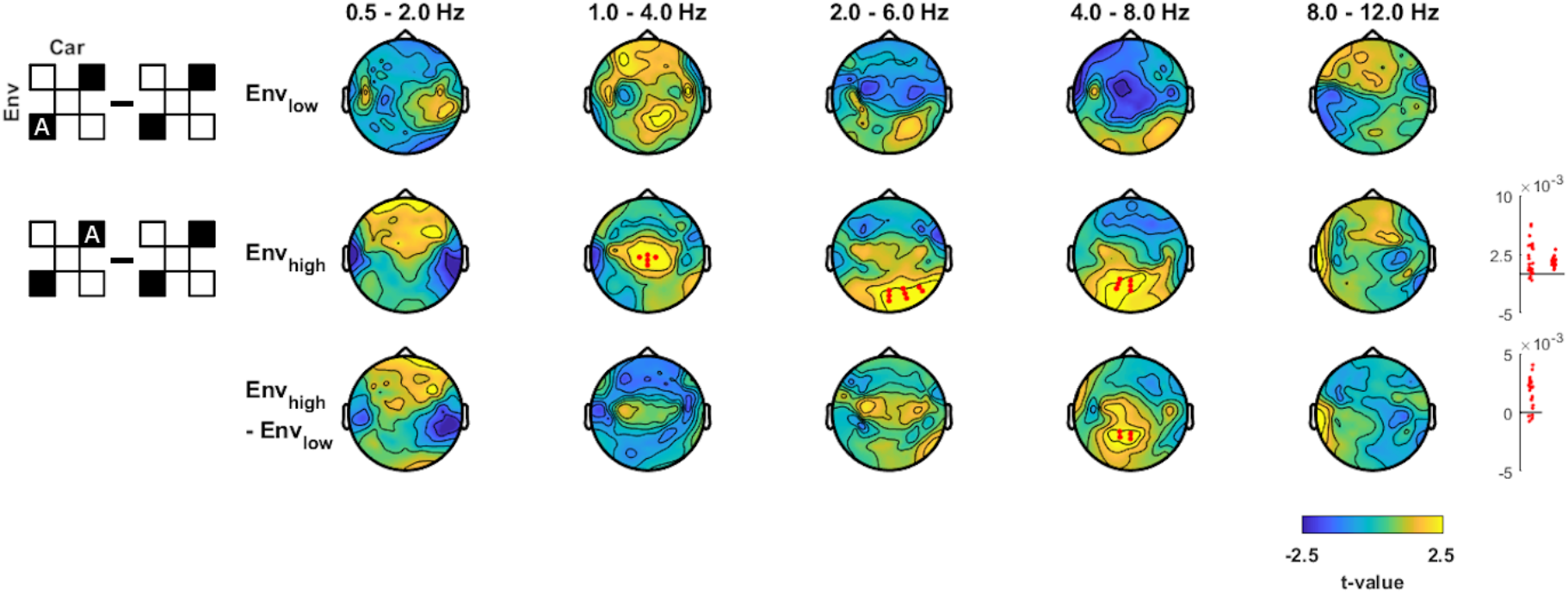
Effect of attention on envelope tracking. Tracking of individual envelopes in conditions where one envelope was actively attended minus conditions in which attention was not focused on a specific envelope. Pictograms on the left indicate which experimental conditions (black filled squares) were contrasted (A indicating the attended signal). Topographies show the electrode-wise t-values for the group-level statistics. Red and white dots mark positive and negative clusters (derived using cluster-based permutation statistics, p<0.01). Panels on the right show MI values for individual participants in each cluster (red for positive and blue for negative clusters).

**Fig. 6.**
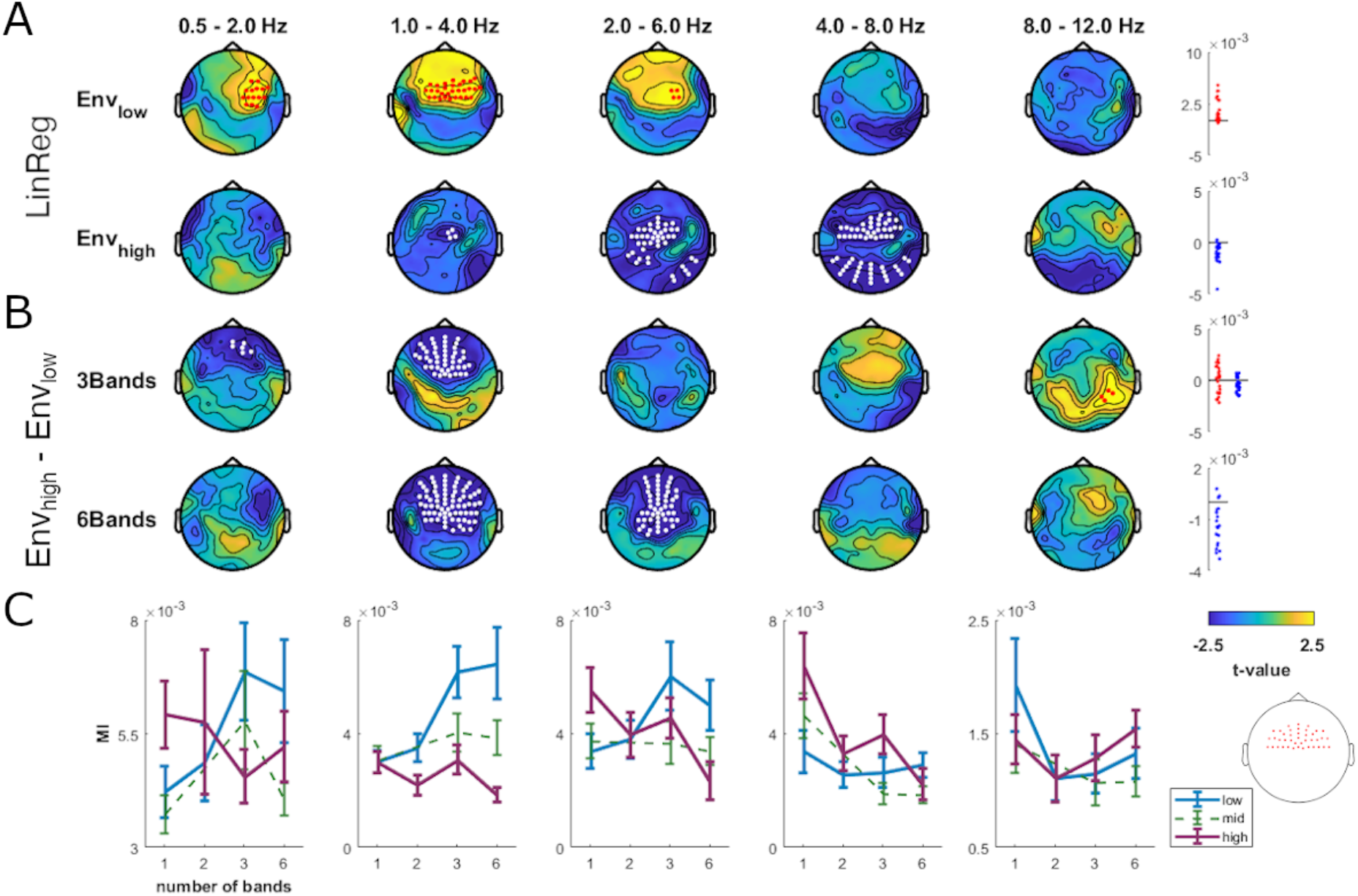
Differential envelope tracking with increased spectral detail. (**A**) Change in envelope tracking across conditions with 1, 2, 3 and 6 concurrently presented envelope-carrier pairs, computed separately for Env_low_ and Env_high_. Regressing MI values (LinReg) for Env_low_ against the number of presented envelope-carrier pairs revealed stronger tracking with more spectral detail in delta1 to theta1 (1 – 6 Hz), while MI values of Env_high_ decreased in delta2 to theta2 (2 – 8 Hz). (**B**) Contrast of MI values for Env_high_ minus Env_low_ in conditions with 3 and 6 simultaneously presented envelope-carrier pairs. Topographies show the electrode-wise t-values for the group-level statistics (derived using cluster-based permutation statistics, p<0.01). Panels on the right show individual data for each cluster (red for positive and blue for negative clusters). (**C**) MI values averaged across fronto-central electrodes (inlay) for Env_low_, Env_mid_, and Env_high_. Error bars indicate s.e.m. across participants.

**Fig. 7.**
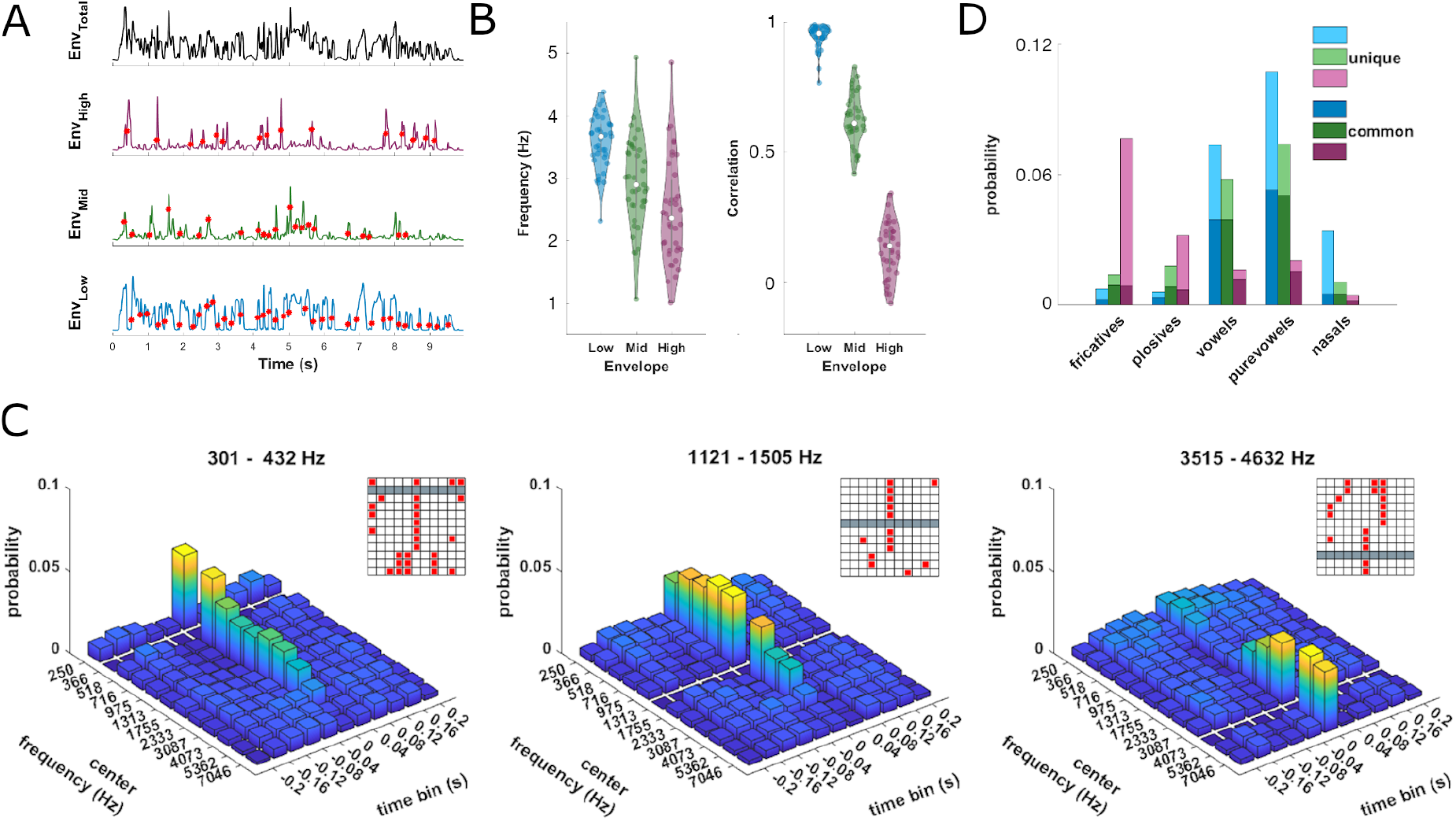
Acoustic features in band-limited speech-derived envelopes. (**A**) Time course of a broadband (0.2 – 8 kHz) and three band-limited envelopes (boundaries between low, mid and high: 0.2, 0.83, 2.66, 8 kHz) from a single sentence. For each envelope band we detected acoustic onsets (red dots) as peaks in the rate of change (first derivative). (**B**) Left: Frequency of onsets in each envelope band across sentences (dots). The lowest band exhibits the highest average onset frequency and is most consistent across all sentences, followed by the middle and the high band. Right: Correlation of band-limited envelopes from three frequency ranges with the overall envelope (see Methods for definition). (**C**) Temporal structure of envelope onsets computed for a finer spectral resolution using 12 bands (see Methods for precise frequency ranges). For each band-limited envelope we determined the joint probability of onsets across bands that fell within a 400 ms window (divided into 40 ms bins). Envelopes are referred to in terms of their center frequency with the respective reference envelope being cleared out (gray bars). Inlays show temporal bins with significant likelihoods (red boxes). (**D**) Phoneme probability near acoustic onsets (+/-40ms), divided by envelope and subgroup. Subgroups are defined as ‘unique’ and ‘common’ phonemes occurring across frequency bands.

## 2. Materials and Methods

### 2.1 Participants and data acquisition

We collected data from 24 native German speaking participants (19 females, mean age = 23.4 ± 3.6 SD). All participants provided written informed consent and were screened for hearing impairment with the speech, spatial and qualities of hearing scale (SSQ) prior to the study (Gatehouse and Noble, 2004). We excluded participants with scores two standard deviations below the mean of a young control group reported by Demeester et al. (Demeester et al., 2012). Participants received monetary compensation of 10 € per hour. The experiment was approved by the local ethics committee of the Bielefeld University, Germany, and was conducted in compliance with the Declaration of Helsinki. Participants were seated in a dimly lit, acoustically and electrically shielded room (Desone Ebox, Germany). Auditory stimuli were presented binaurally using Sennheiser headphones (Model HD200) from a Creative sound blaster z Soundcard at an average intensity of 68 dB SPL. EEG signals were continuously recorded using an active 128 channel BioSemi system using Ag-AgCl electrodes mounted on an elastic cap (BioSemi). Four additional electrodes were placed at the outer canthi and below the eyes to obtain the electrooculogram. Electrode impedance was kept at <40 kΩ. Data were acquired at a sampling rate of 1024 Hz using a lowpass filter of 400 Hz.

### 2.2 Speech material and stimuli

The original material based on which the experimental stimuli were modelled comprised 39 sentences extracted from German radio interviews conducted by the Deutschlandfunk on various topics. The sentences were spoken by nine different professional speakers (2 females, 7 male) and lasted on average 8.65 ± 2.3 s (mean ± SD, ranging from 4.13 s to 13.67 s) and comprised in total 337.5 s of spoken material. The sentences consisted of 22.1 ± 7.2 words. All recordings are publicly available online and were downloaded and converted to .wav files with a sampling rate of 44.1 kHz (https://www.deutschlandradio.de/audio-archiv.260.de.html).

To create the experimental stimuli, we first extracted amplitude-envelopes from the spoken sentences and then applied these to bandpass filtered Gaussian white noise with varying center frequencies as described below. The band-limited envelopes were obtained by bandpass filtering each sentence into logarithmically spaced frequency bands covering the range from 0.2 to 8 kHz. Depending on the experimental condition or analysis, we used either three frequency bands (boundaries: 0.2, 0.83, 2.66, 8 kHz), six bands (boundaries: 0.2, 0.43, 0.83, 1.5, 2.66, 4.6, 8 kHz) or twelve bands (boundaries: 0.2, 0.3, 0.43, 0.6, 0.83, 1.1, 1.5, 2, 2.66, 3.5, 4.6, 6.1, 8 kHz). In the following we often refer to low, mid, and high bands, which here are defined as follows: low: 0.2 – 0.83 kHz, mid: 0.83 – 2.66 kHz and high: 2.66 – 8 kHz. For filtering we applied a zero-phase 4th order Butterworth IIR filter using the Matlab filtfilt function. Band-limited envelopes were computed by taking the absolute of the Hilbert transform (Fig. 1A). To construct stimuli, we bandpass filtered Gaussian white noise into the same logarithmically spaced frequency bands and applied the envelopes extracted from the speech material to create amplitude-modulated band-limited noises. Similar to the envelopes, carriers are thus defined by their respective frequency bands (e.g. low, mid and high). To create different experimental conditions, we constructed stimuli by combining these band-limited envelopes not only with a noise carrier at the same frequency (i.e. ‘natural’, Fig. 1B, signals 1 and 5), but also by applying the envelope of one frequency band to a different carrier band (i.e. ‘unnatural’, Fig. 1B, signals 2 and 4). We also constructed multiband stimuli by presenting 2, 3, 6, or 12 carriers simultaneously, each modulated by the same or distinct envelopes, as described further below.

### 2.3 Experimental design

The experiment consisted of five blocks, in which participants were presented with different combinations of amplitude-modulated noises. All blocks were presented within the same session in a fixed order. During the entire experiment, participants were instructed to fixate a central fixation cross and to blink as little as possible. Envelopes derived from all 39 sentences were repeated in a pseudorandom sequence in each experimental condition. The first block comprised five conditions, in each of which we presented one carrier modulated with one envelope, as depicted in figure 1B. Three conditions presented natural combinations of carrier frequency and band-limited envelope, e.g. Env_low_ with Car_low_ (Fig. 1B, signals 1, 3 and 5), two conditions presented unnatural pairings, i.e. Env_low_ with Car_high_ and Env_high_ with Car_low_ (signals 2 and 4). In the second block we presented a mixture of two amplitude-modulated noises, whereby the two carriers at different frequencies were modulated with the same bandlimited envelope: one condition presented Env_low_ modulated on Car_low_ together with Env_low_ modulated on Car_high_ (i.e. signals 1 plus 2), and one condition presented Env_high_ modulated on Car_low_ together with Env_high_ modulated on Car_high_ (i.e. signals 4 plus 5). In the third and fourth blocks we presented a mixture of two amplitude-modulated noises, whereby each noise was modulated by its natural envelope (i.e. Env_low_ with Car_low_ and Env_high_ with Car_high_; Fig. 1B), while also manipulating participants’ attentional demands. In block three participants were asked to attend both carriers, while in block 4 they were asked to attend either the low or the high carrier. To ensure attentive engagement in blocks 1 to 4, participants performed a dummy task in which they had to report the occurrence of a silent gap within the stimulus. These gaps were randomly distributed over all trials and occurred in 20 % of trials. Each gap lasted between 450 and 600 ms and randomly occurred within the third quarter of the stimulus. In block three, participants were tasked to detect a gap which was implemented to appear simultaneously in both streams, whereas in block four participants were tasked to attend either the low or the high carrier and only respond to a gap occurrence in that attended carrier.

In the fifth block participants were presented with more than two simultaneous carriers, similar to noise-vocoded speech. Each sentence was repeated in each of three conditions, comprising either 3, 6, or 12 envelope-carrier pairs. For the 3-band condition, carriers and envelopes were derived from the same three bands (low, mid, high) as used in Blocks 1 – 4. For the 6 and 12 bands, carriers and envelopes were derived from more narrow bands (see filtering procedures above). Participants were asked to pay attention in order to perform a comprehension task. Subsequent to each sentence they had to report which out of three candidate words they had heard by pressing one of three buttons. Additionally, they were asked to rate their confidence in their decision on a scale from 1 (low) over 2 (medium) to 3 (high). As each sentence was repeated in each sub-block, this introduced a potential confound of memory and recognition. Given that performance was already high for 6 bands, we did not investigate the subsequently presented 12-band condition any further, as the cerebral encoding and perceptual reports may be confounded by memory-related effects.

### 2.4 EEG preprocessing

Data analysis was performed offline using the FieldTrip toolbox (Oostenveld et al., 2011) Version 20190905 and custom written MATLAB scripts. Data and code used in this study will be made available on the Data Server of the University of Bielefeld upon acceptance of the article. The EEG data were bandpass filtered between 0.2 and 60 Hz using a zero-phase 4th order Butterworth IIR filter and resampled to 150 Hz. To detect channels with excessive noise we selected channels with an amplitude above 3.5 standard deviations from the mean of all channels and interpolated these using a weighted neighbor approach: on average we interpolated 13.55 ± 6.73 channels across subjects (mean ± SD). Noise cleaning was performed using independent component analysis based on 40 components across all blocks simultaneously. Artifacts were detected automatically using predefined noise templates capturing typical artifacts related to eye movements, blinks or localized muscle activity based on our previous work (Kayser et al., 2016; McNair et al., 2019). On average 12.16 ± 4.76 components were removed (mean ± SD). Finally, all channels were referenced to the mean of eight temporal electrodes (B14, B25, B26, B27, D8, D22, D23, D24).

### 2.5 EEG analysis

To quantify the relationship between the EEG and the stimulus envelope we used a mutual information (MI) approach (Ince et al., 2017). For each trial we omitted the first 500 ms to remove the evoked onset response and then separately computed the MI for multiple lags between stimulus and EEG, ranging from 60 to 140 ms in 20 ms steps (Keitel et al., 2018). We subsequently averaged the MI values over all lags. To compute the MI values, the single trial EEG data was cut to the same length as the corresponding acoustic envelope. Then, we appended all trial-wise envelopes and EEG data from the same condition and filtered these arrays into partly overlapping frequency bands: delta1 (0.5 – 2 Hz), delta2 (1 – 4 Hz), theta1 (2 – 6 Hz), theta2 (4 – 8 Hz) and alpha (8 – 12 Hz), respectively using a zerophase 4th order Butterworth IIR filter. To compute the mutual information between EEG and stimulus vectors we derived the analytic signal using the Hilbert transform and calculated the MI using the Gaussian copula approach (Ince et al., 2017). By using the complex signals of the Hilbert transform we effectively quantified the relationship between two signals using both power and phase information. Analysis in block 5 of the experiment was conducted by computing the MI between the EEG signal and the three envelopes (i.e. Env_low_, Env_mid_, Env_high_) for both the 3 and 6 envelopes condition. Specifically, in the 6 envelope condition, we first averaged the two envelopes covering the low, mid or high ranges respectively and then computed the MI between these averaged envelopes and the EEG. This enabled us to compare tracking of equal spectral distances across conditions with varying detail.

### 2.6 Statistical analysis

For statistical analysis of the EEG data we relied on a cluster-based permutation approach, relying on 2000 permutations, requiring a minimum of three significant neighboring electrodes and using the ‘cluster-mass’ as cluster-forming criterion (Maris and Oostenveld, 2007). To test whether the tracking of envelope-noise pairs in the EEG was significant when compared to a condition of no systematic relation between stimulus and brain activity (Fig. 2), we computed a distribution of randomized values obtained by computing the MI between EEG and time shifted speech envelopes 2000 times. For this, all trial-wise appended stimulus arrays were circularly shifted by a uniformly distributed random lag between ¼ N and N – ¼ N, where N is the number of samples in the stimulus array. By this we ensured that the stimuli were in no relation to the brain signal, while still maintaining their spatially and spectrally distinct autocorrelation. We then formed clusters based on electrode-frequency bins exceeding a cluster-forming threshold of p<0.01, i.e. exceeding the 99^th^ percentile of the frequency-specific randomized distribution. Frequency-specific randomization was used as the MI values decrease as a function of the chosen pass-band frequency and therefore the bias cannot be assumed to be drawn from the same null distribution. Clusters were considered significant if they exceeded a two-sided alpha level of 0.01.

To compare the MI between conditions, we first computed electrode- and frequency-wise paired two-tailed t-tests across participants. Electrodes were considered if their t-value exceeded a critical level of p<0.01. Then, to create a null distribution under the hypothesis of no difference, we randomly flipped the sign of the conditional MI differences to obtain a randomization distribution (Maris and Oostenveld, 2007). To control for multiple comparisons across electrodes and EEG frequency bands the maximum statistics was used. In cases where nearby clusters reflected a similar effect but were separated by at most one electrode, we conjugated these into one cluster. Instances of such conjugations are indicated in the text. For each significant cluster we computed the sum of within-cluster values as a cluster statistic (cstat). Additionally, the electrode with the maximum (for positive or minimum for negative clusters) test statistic was selected and subsequently used to compute Cohen’s d as an indicator of effect size. To investigate changes in speech tracking with the number of envelope bands (Fig. 6), we regressed the MI values for each electrode and frequency band against the number of presented envelope-carrier pairs. We applied cluster-based permutation to test regression betas against zero.

The spatial similarity of MI topographies was obtained as the Pearson correlation coefficient between the group-averaged topographies in each frequency band, averaged across envelopes and carriers. Statistical significance of these correlations was tested by obtaining group-level bootstrap confidence intervals, created by randomly sampling participants with replacement 2000 times.

To test for differences in the frequency of acoustic onsets between envelope bands we compared their onset frequency with a one-way repeated measures ANOVA followed by a post hoc Tukey-Kramer multiple comparison test. Additionally, to test the joint onset probability for significance we created a null distribution by randomly shifting each envelope separately in time and computing the randomized likelihood 2000 times. For all statistical tests we provide exact p-values, except where these are below 10^-3^.

### 2.7 Analysis of stimulus material

We analyzed statistical regularities of the acoustic onsets in the speech material as these provide prominent landmarks that drive the neural entrainment to speech (Drennan and Lalor, 2019; Khalighinejad et al., 2017; Oganian and Chang, 2019). For each band-limited envelope we computed its derivative and determined local peaks using Matlab’s ‘findpeaks’ function with a minimum distance of 125 ms between neighboring peaks. For each sentence we then computed the onset frequency by dividing the number of onsets by the length of the given sentence. This was done for each envelope band separately and subsequently averaged over sentences. Additionally, we computed the correlation between each band-limited envelope band and the overall envelope, which was defined as the average of the all band-limited three envelopes derived from band-limited speech (Fig. 7A – B). To determine whether onsets in different bands were temporally structured we determined their joint probability: for each onset in a given envelope band we calculated the probability of onsets in all remaining bands that fell within a 400 ms window using temporal bins of 40 ms length (Fig. 7C).

To link the acoustic onsets with their phonemic content, we examined how these onsets coincide with phonemes. For this we used the web-based WebMAUS Basic tool by the Bavarian Archive for Speech Signals to force-align written transcripts to the acoustic waveform (Kisler et al., 2017). Phonemic onsets were stored as TextGrid files and subsequently adjusted by hand using Praat (Boersma and van Heuven, 2001). For each envelope band we determined the closest phoneme onset to each acoustic onset within a temporal distance of 40 ms. Phoneme counts were grouped by their manner of articulation (fricatives, plosives, vowels, pure vowels, nasals) (Fig. 7D). Phoneme counts for each envelope were then divided into two subgroups: ‘unique’ phonemes that coincide with acoustic onsets in only one band, and ‘common’ phonemes coinciding with acoustic onsets in more than one envelope band. To compute the probability of phoneme groups given any acoustic onset, we normalized the count of each phoneme group to the total number of phonemes in all sentences.

## 3. Results

Across five blocks participants listened to acoustic stimuli presenting band-limited noises modulated by speech-derived band-limited envelopes. Within and between blocks we manipulated the spectral match between envelope (i.e. the frequency range in the natural speech from which the envelope was extracted) and carrier (i.e. the spectral band of the respective noise), participants’ attention to specific envelope-carrier pairs, and varied the number of simultaneously presented envelope modulated carriers.

### 3.1 Behavioral performance

In the first three blocks participants were asked to detect a silent gap to ensure they paid attention to the stimuli. Gaps were reported with an accuracy of 95.4 ± 0.4 % (mean ± SD; averaged over participants and blocks 1 – 3). In the fourth block, participants were presented with two carriers but had to detect a gap in one ‘attended’ stimulus: they did so correctly in 79.3 ± 9.1 % of trials. In the fifth block, participants listened to noise-vocoded speech with varying spectral detail (3, 6 and 12 envelopes) and were asked to perform a 3-choice word recognition task. Accuracy increased from 45.1 ± 11.4 % to 91.8 ± 6.7 % and 97.9 ± 4.0 % across conditions. In this block they also indicated their confidence on an interval scale from 1 to 3, which increased over the three conditions from 1.34 ± 0.43 to 2.65 ± 0.21 and 2.93 ± 0.08. Because in Block 5 performance already reached near ceiling level for 6 bands, we omitted the 12-band condition from the subsequent analysis, as this may be confounded by recognition or memory effects.

### 3.2 Tracking of spectrally limited speech envelopes

In block 1 we presented five conditions each presenting one carrier band modulated by one speech-derived narrow band-limited envelope, while varying the spectral match between carrier and envelope bands. Three conditions presented natural pairings, i.e. the envelope and carrier were derived from the same frequency range, while two conditions presented unnatural pairings (Fig. 1B). For each condition we quantified the envelope tracking as the mutual information between the complex-valued representation of the envelope and the EEG activity (Ince et al., 2017; Keitel et al., 2017).

First, we quantified envelope tracking separately for carrier noises and for envelopes derived from the low and high frequency bands, averaging over the respective other dimension (e.g. quantifying the tracking when envelopes were presented at Car_low_ regardless of which envelope was used Fig. 2). For each of these four analyses, tracking prevailed in lower EEG bands and over central and temporal electrodes, as expected from previous work (Giordano et al., 2017; Kayser et al., 2015; Keitel et al., 2018). To ensure that these tracking signatures were indeed robust, we contrasted the actual MI values to a null-distribution obtained under the assumption of no reliable association between stimulus and EEG signal. For each envelope and each carrier band, all electrodes formed one single significant cluster over all frequency ranges (cluster-based permutation test; all p<10^-3^, Car_low_: cstat=2.03, Car_high_: cstat=2.15, Env_low_: cstat=1.85, Env_high_: cstat=2.44).

We then asked whether the neurophysiological processes giving rise to these MI topographies across analysis factors share a similar spatial layout, or rather reflect spatially distinct generators. The similarities between condition-averaged MI topographies were significantly correlated across neighboring EEG bands (Fig. 2B, group-level bootstrap test, p<10^-3^), revealing significant similarities between each neighboring topographies (0.5 – 12 Hz) and additionally between delta2 and theta2 (0.5 – 2 Hz and 4 – 8 Hz). Concerning the role of carrier or envelope bands, we found that the topographies obtained using different carriers were significantly correlated (averaged correlation across EEG bands; r=0.88, group-level bootstrap test p<10^-3^), while the topographies obtained using low and high frequency envelopes were only weakly but still significantly correlated (r=0.29, p<10^-3^). These results suggest that envelope tracking in different bands of the EEG likely reflects distinct neurophysiological processes. These tracking signatures are not influenced by the choice of carrier frequency while acoustic envelopes extracted from low (0.2 – 0.83 kHz) and high (2.66 – 8 kHz) spectral ranges seem to be reflected by distinct neurophysiological signatures.

### 3.3 High-frequency envelopes are reflected more strongly in the EEG

Addressing the first of our main questions, we asked whether the signatures of envelope tracking are specific to the match between the carrier frequency and the spectral range from which the envelope was extracted. A direct contrast of MI values between conditions in which Car_low_ or Car_high_ were presented (Fig. 3, ΔCar) revealed no significant effects except in the alpha band, where MI values were higher for Car_high_ (p<10^-3^, cstat=-20.9, cohen’s d=-0.93). A direct contrast between conditions in which either Env_low_ or Env_high_ were presented (Fig. 3, ΔEnv) revealed higher MI for Env_high_ over central channels in delta1 (0.5 – 2 Hz; p<10^-3^, cstat=52.51, cohen’s d=0.91), over fronto-central channels in theta1 and theta2 (2 – 8 Hz; p<10^-3^, cstat=320.37, cohen’s d=1.21), being conjugated from two separate clusters, and over occipital channels in the same frequency band (2 – 8 Hz; p<10^-3^, cstat=97.66, cohen’s d=0.88). Finally, we asked whether there is an interaction between envelope and carrier bands. For this we compared unnatural (Fig 1B, signals 2 plus 4) and natural pairings (signals 1 plus 5) (Fig. 3, Inter), and found no significant difference. Together with the topographical differences in the tracking of Env_low_ and Env_high_ (c.f. Fig. 2), these results suggest that speech envelopes derived from lower and higher spectral ranges are reflected differently in the EEG.

### 3.4 Contextual information influences tracking of Env_high_ more than Env_low_

Using the conditions in blocks 2 and 3 we examined whether and how the tracking of individual envelopes is influenced by concurrent information presented to the listener. First, we asked whether the tracking of a specific envelope is enhanced when the same acoustic envelope is presented in a broader spectral range: In block 2, we presented carriers at two frequency ranges, each modulated by the same envelope (2Car condition; Fig. 3). Comparing this 2-carrier condition to the respective 1-carrier condition (from block 1) revealed higher MI values for the 2-carrier condition: for Env_low_ over right posterior electrodes in theta1 (Fig. 3, 2 – 6 Hz; Env_low_^2Car^, p<10^-3^, cstat=11.08, cohen’s d=0.63); for Env_high_ over right posterior electrodes in delta 2 (1 – 4 Hz; p<10^-3^, cstat=64.19, cohen’s d=0.81) and over left posterior electrodes in theta2 (4 – 8 Hz; p<10^-3^, cstat=33.50, cohen’s d=0.80). This shows that envelope tracking is enhanced when the same information is present in a broader spectral range.

In block 3 we presented two carriers each modulated by their respective matching (natural) envelopes in order to probe whether the tracking of a specific envelope is enhanced when the complementary speech-like information is presented in another spectral range (i.e. presenting Env_low_ carried by Car_low_ and Env_high_ carried by Car_high_ together Fig. 4). MI values were lower in the 2-carrier condition compared to the respective values obtained from block 1: for Env_low_ MI was reduced over right central-posterior electrodes in theta2 (Fig. 4, 4 – 8 Hz; p<10^-3^, cstat=-28.16, cohen’s d=-0.78); for Env_high_ MI values were reduced over posterior electrodes in the theta1 to alpha bands (2 – 12 Hz; p<10^-3^, cstat=-232.82, cohen’s d=-1.04) and over central electrodes in theta2 band (4 – 8 Hz; p<10^-3^, cstat=-46.02, cohen’s d=-0.66). A direct comparison between Env_high_ and Env_low_ revealed a stronger reduction of MI values for Env_high_ over central channels in delta2 to theta1 (1 – 6 Hz; p<^10-^3, cstat=-50.62, cohen’s d=-0.69) and over left posterior sites in theta2 (4 – 8 Hz; p<10^-3^, cstat=62.18, cohen’s d=-0.99). Hence, the tracking of a specific envelope-carrier pair in the theta band becomes less prominent when an additional complementary speech envelope is presented and this effect is particularly prominent for Env_high_.

### 3.5 Attention influences tracking of Env_high_ more than Env_low_

We then asked how focused attention affects the tracking of a specific envelope. In block 4 we presented a mixture of two natural envelope-carrier pairs similar to block 3, but asked participants to attend to either Env_low_ or Env_high_. The contrast between the unfocused attention in block 3 (target gap in both envelopes) and block 4 (focused attention) revealed no effect for Env_low_, while the MI for Env_high_ was enhanced by attention over central electrodes in delta2 (Fig. 5, 1 – 4 Hz; p<10^-3^, cstat=13.13, cohen’s d=0.57) and over posterior electrodes in theta1 and theta2 (2 – 8 Hz; p<10^-3^, cstat=42.14, cohen’s d=0.68). To understand whether the attentional modulation differed between Env_low_ or Env_high_, we directly contrasted the attention effect between these (Fig. 5, Env_high_ – Env_low_). This revealed that attention had a larger influence on the tracking of Env_high_ than of Env_low_ over central posterior electrodes in theta2 (4 – 8 Hz; p<10^-3^, cstat=11.44, cohen’s d=0.63).

### 3.6 Tracking of Env_low_ prevails when the acoustic signal becomes intelligible

Combining the data from blocks 1, 2 and 5 we asked how the tracking of one specific band-limited envelope presented among a mixture of multiple natural envelope-carrier pairs is affected when the amount of overall presented spectral detail is increased. For this analysis, we considered from the first two blocks only the natural envelope-carrier pairs, which presented either 1 or 2 natural envelope-carrier pairs simultaneously. From block 5 we analyzed the conditions presenting 3 or 6 simultaneous naturally-matched envelope-carrier pairs.

First, we investigated how the tracking of Env_low_ and Env_high_ scaled with the number of simultaneously presented envelope-carrier pairs (1, 2, 3, or 6 pairs; Fig. 6A). The MI values for Env_low_ generally increased: in delta1 to theta1 over fronto-central electrodes (linear regression, LinReg, of MI against the number of carriers: 0.5 – 6 Hz; p<10^-3^, cstat=150.68, cohen’s d=0.84). In contrast, MI values for Env_high_ generally decreased in delta2 to theta2 over central and posterior electrodes (1 – 4 Hz; p<10^-3^, cstat=-342.21, cohen’s d=-0.96). To confirm this differential effect, we performed a direct contrast between Env_low_ and Env_high_ for each number of presented bands (Fig. 6B). For the 3-band condition, which was generally unintelligible (Fig. 6B), we found higher MI for Env_high_ in alpha over right central parietal electrodes (8 – 12 Hz; p<10^-3^, cstat=11.71, cohen’s d=0.64) and higher MI for Env_low_ in delta1 and delta2 over fronto-central electrodes (0.5 – 4 Hz; p<10^-3^, cstat=-154.0, cohen’s d=-0.70). For 6 bands, when participants could readily understand the sentences, we found stronger tracking of Env_low_ in a cluster ranging from delta2 to theta1 over fronto-central electrodes (1 – 4 Hz; p<10^-3^, cstat=-380.12, cohen’s d=-0.84). Figure 6C directly illustrates this differential tracking of Env_high_ compared to Env_low_ over fronto-central channels in the 1-6 Hz EEG when an increasing number of envelope-carrier pairs is presented.

### 3.7 Statistical properties of band-limited envelopes

To understand the temporal structure of the band-limited speech envelopes we investigated their statistical regularities (Fig. 7A). First, we compared the similarity (Pearson correlation) between each band-limited envelope and the overall envelope (Fig. 7B). While Env_low_ (0.2 – 0.83 kHz) was strongly correlated with the overall envelope (R^2^=0.89 ± 0.01, mean ± s.e.m. across sentences), the similarities between Env_mid_ (0.83 – 2.66 kHz) and Env_high_ (2.66 – 8 kHz) and the overall envelope were much reduced: a one-way ANOVA revealed a significant effect of band (F(2,114)=6.57, p<10^-4^, η^2^=0.94) and a Tukey-Kramer post hoc comparison revealed all pairwise comparisons as significant (each p<10^-4^). Second, we investigated the structure of acoustic onsets (Fig. 7A, dots). Focusing on these three spectral bands we found that Env_low_ featured the highest onset frequency (3.57 ± 0.44 Hz), followed by Env_mid_ (2.94 ± 0.73 Hz) and Env_high_ (2.46 ± 0.83 Hz; Fig 7B). A one-way ANOVA revealed a significant difference between bands (F(2,114)=26.32, p<10^-4^, η^2^=0.32) and a Tukey-Kramer post hoc comparison revealed all pairwise comparisons as significant (each p<0.0065). We then asked if acoustic onsets are temporally co-structured between bands. For this, we computed their joint probability across frequency bands and time-lags, using a division into 12 bands (Fig. 7C). This revealed that onsets in neighboring bands occur at the same time, but this dependency decreases distinctively with spectral distance: acoustic onsets in bands higher than about 3 kHz were largely independent from onsets in bands below 3 kHz.

Last, we asked whether the acoustic onsets are related to the phonemic content of each envelope. For this we computed the probability of occurrence of individual phonemes in the original speech material near the acoustic onsets in band-limited envelopes and divided these occurrences into those ‘unique’ and ‘common’ between bands (Fig. 7D). Acoustic onsets in Env_high_ coincided most with fricatives and plosives and were largely unique to these. On the contrary, acoustic onsets from Env_mid_ and Env_low_ were more likely to coincide with vowels, pure vowels and nasals. Together these results show that speech envelopes derived from distinct frequency ranges are temporally independent and carry partially complementary information about the occurrence of distinct types of phonemes.

## 4. Discussion

The representation of speech in the brain is often examined by measuring the alignment of rhythmic brain activity to the acoustic envelope of the signal (Ahissar et al., 2001; Ding et al., 2014; Giraud and Poeppel, 2012; Gross et al., 2013; Kayser et al., 2015; Oganian and Chang, 2019; Teng et al., 2019; Teoh et al., 2019). To quantify this alignment many studies, rely on the overall or broadband acoustic envelope, which describes the amplitude fluctuation of the signal across the full spectral range and which provides a convenient and low-dimensional representation for data analysis. Our results show that this overall envelope combines acoustic signatures that are seemingly encoded by distinct neurophysiological processes giving rise to spatially distinct signatures of acoustic tracking in the human EEG. Because low and high frequency speech-derived envelopes also relate to separate acoustically and phonetically features (Fig. 7), the neural processes encoding these spectrally distinct envelopes likely carry complementary information relevant to acoustic and speech-specific information.

### 4.1 The overall envelope provides a distorted picture of speech tracking

We probed how synthetic and incomprehensible sounds carrying typical regularities of natural speech-derived envelopes are reflected in low frequency EEG activity. Specifically, we asked whether these are represented in a similar manner as real speech. In large our data show that this is indeed the case: Envelopes were tracked over fronto-central electrodes at EEG frequencies below 8 Hz, consistent with previous studies (Drennan and Lalor, 2019; Etard and Reichenbach, 2019; Kayser et al., 2015; Mai and Wang, 2019; Synigal et al., 2019). However, the envelopes derived from a higher spectral range (defined here as 2.66 – 8 kHz) were reflected in spatially distinct EEG topographies compared to the envelopes from the lower spectral range (i.e. 0.2 – 0.83 kHz), suggesting that spectrally segregated acoustic envelope features are reflected in segregated neurophysiological processes (Fig. 2).

Interestingly, the differential tracking of Env_high_ and Env_low_ was not specific to the frequency of the respective carrier noise, indicating that the underlying processes are not selective to the natural spectral match of carrier and speech-like envelope (Fig. 2), at least for the synthetic sounds used here. In how far this interaction between envelopes complies to intelligible speech has still to be investigated. Together with the differential sensitivity of the tracking of Env_high_ and Env_low_ to contextual and attention manipulations (Fig. 3,4), this implies that combining band-limited envelopes into an overall envelope for data analysis provides a distorted picture on the encoding of complex sounds in dynamic brain activity: both in terms of the topographical and spectral distribution of the respective neurophysiological signatures, and in terms of their sensitivity to experimental manipulations. Given that the representation of Env_low_ prevailed for rich and comprehensible sounds (e.g. when presenting 6 bands simultaneously; Fig. 6), and given the high correlation between Env_low_ and the overall envelope (Fig. 7), we posit that studies focusing on the overall speech envelope mostly capture the tracking of the speech envelope from the lower spectrum. Yet, when the acoustic signal is impoverished this could confound distinct neurophysiological processes.

### 4.2 Differential encoding of low and high frequency-derived envelopes

Our data show that the envelopes obtained from low and high frequency ranges are reflected differentially in the 1 – 6 Hz EEG. The tracking of naturalistic sounds in dynamic brain activity has been frequently related to the neural encoding of acoustic onsets, in particular as acoustic transients drive individual neurons and induce an alignment of rhythmic brain activity to the acoustic signal (Daube et al., 2019; Drennan and Lalor, 2019; Khalighinejad et al., 2017; Oganian and Chang, 2019). Such transients are not exclusive to natural speech but are similarly present in band-limited noises. Our results demonstrate that the neurophysiological process reflecting such band-limited envelopes are largely similar, or even the same, as those engaged by real speech. Hence, the typical tracking signatures investigated in studies using speech as a stimulus are not exclusive to ecologically meaningful stimuli. At the same time, our data also suggest that the preferential tracking of the low frequency envelope for (near) intelligible speech may be functionally meaningful. The analysis of the acoustic speech material showed that low and high frequency envelopes can reflect independent acoustic landmarks which relate, in part, to the unique signaling of the occurrence of specific phoneme groups. The low frequency envelope in real speech carries more energy (Peelle et al., 2013), and lowpass filtered speech has been shown to be sufficient for comprehension (Elliott and Theunissen, 2009). Prominent acoustic landmarks in the low frequency envelope relate to the occurrence of a large group of phonemes, and hence the prominent tracking of the low frequency envelope may reflect an adaptation to the typically most robust aspect of ecologically relevant sounds such as speech.

Several previous studies have exploited narrow-band speech envelopes to study the relation between stimulus and brain activity, for example by exploiting this detailed acoustic information to boost the performance of forward models predicting the neural response from multiple stimulus dimensions (Daube et al., 2019; Di Liberto et al., 2015; Ding and Simon, 2012). For example, models based on spectrally refined envelope signatures (i.e. the stimulus spectrogram) outperformed phoneme-based models for high frequency EEG bands such as alpha, whereas the phoneme-based models yielded higher prediction accuracies for delta and theta band EEG signals (Di Liberto et al., 2015). Related studies also showed that low and high-frequency derived envelopes are seemingly reflected with distinct latencies in the EEG or differentially affected by attention (Brodbeck et al., 2019; Di Liberto et al., 2015). However, these previous studies did not directly quantify or contrast such spectrally-limited effects, but merely indicated that distinct acoustic features are possibly represented by separate neural processes. The present work directly addresses these questions and provides a systematic and quantitative assessment how narrow band speech-derived envelopes are reflected in dynamic brain activity.

### 4.3 The neurophysiological processes underlying envelope tracking

Our results corroborate the notion that the tracking of acoustic envelopes in delta and theta band brain signals represent functionally distinct processes. Delta-band tracking (below 4 Hz) has been shown to covary with experimental manipulations affecting intelligibility (Ding et al., 2016; Zion Golumbic et al., 2013; Zoefel et al., 2018) and has been related to temporally extended supra-syllabic features (Keitel et al., 2018; Mai and Wang, 2019). Nevertheless, it is still debated whether delta tracking is causally related to comprehension. While some studies advocate for such a role (Etard and Reichenbach, 2019; Peelle et al., 2013; Wilsch et al., 2018; Zoefel et al., 2018), experiments using pop-out stimuli did not find a clear relation (Baltzell et al., 2017; Millman et al., 2015) and a manipulation of speech rhythm resulted in reduced delta tracking while preserving intelligibility (Kayser et al., 2015). In the present data tracking of Env_low_ in the 1 – 4 Hz EEG increased monotonically with acoustic detail rather than correlating with comprehension performance, comparable to recent findings that demonstrate increasing envelope representation with the number of envelope bands (Prinsloo and Lalor, 2020). One possibility is that delta band tracking reflects processes that screen sounds for speech-like regularities, such as lexical structures or prosody, to which early stage spectro-temporal filters could be flexibly tuned to (Brodbeck et al., 2019; Khalighinejad et al., 2019; Schönwiesner and Zatorre, 2009). These processes then facilitate neural encoding prior to comprehension (Giraud and Poeppel, 2012; Keitel et al., 2018; Schroeder and Lakatos, 2009; Scott, 2019).

Theta band tracking has been linked to the bottom up processing of the acoustic input, in particular of syllabic and sub-syllabic features (Ding et al., 2014; Ding and Simon, 2013; Etard and Reichenbach, 2019; Keitel et al., 2018; Mai and Wang, 2019; Peelle et al., 2013; Rimmele et al., 2015; Scott, 2019). In contrast to some previous studies (Ding et al., 2014; Peelle et al., 2013; Rimmele et al., 2015), in the present data tracking in theta (4 – 8 Hz) and alpha bands (8 – 12 Hz) did not increase with detail. Based on a similar observation a recent study argued that stronger theta tracking of impoverished sounds may reflect effects of attention induced by increased listening effort (Hauswald et al., 2019). However, the apparent discrepancies between studies may also arise from the distinct approaches used to study speech tracking: while some studies used encoding or reconstruction methods (Daube et al., 2019; Hausfeld et al., 2018; Zion Golumbic et al., 2013), others quantified the temporal alignment of speech envelope and brain activity (Hauswald et al., 2019; Peelle et al., 2013), and yet others quantified the between-trial alignment of brain activity (Ding et al., 2014; Rimmele et al., 2015). Furthermore, for vocoded speech or speech-in-noise some studies considered the envelope of the original speech (Ding and Simon, 2013; Peelle et al., 2013) while others used the envelope of the actually presented sound (Hauswald et al., 2019; Millman et al., 2015), possibly contributing to discrepancies in previous work. In the present study, we quantified tracking of the same band-limited envelopes across conditions. With this approach theta band tracking did not exhibit a systematic in- or decrease with the number of simultaneous envelopes presented, but tracking of Env_high_ was significantly reduced, further endorsing the evidence that theta tracking is sensitive to attentional and contextual manipulations (Hauswald et al., 2019). However, a careful assessment of how different methodological approaches to quantify speech tracking affect the specific results would certainly be helpful to resolve some controversies in the previous literature.

Where are the neurophysiological processes giving rise to these tracking signatures located? Early auditory regions entrain to acoustic regularities (Ding and Simon, 2014; Lakatos et al., 2019; Meyer, 2018; Obleser and Kayser, 2019) but do so spatially specific along the tonotopic axis (Lakatos et al., 2013; O’connell et al., 2014). If these regions gave rise to the observed tracking signatures, one should expect a sensitivity of these to the match between the carrier frequency and the spectral range at which the envelope is extracted. However, we found no evidence for this in the delta and theta bands typically associated with acoustic envelope tracking, suggesting that these differential tracking signatures between low and high frequency envelopes arise from higher superior temporal regions that encode temporal and spectral information largely independently (Sohoglu et al., 2020). Indeed, previous neuroimaging studies and direct intracranial recordings showed that delta and theta tracking prevails in large parts of the temporal lobe, but most clearly establish intracortical signals of speech entrainment along superior temporal regions (Gross et al., 2013; Hamilton et al., 2020; Keitel et al., 2018; Mégevand et al., 2020; Moses et al., 2019; Peelle and Davis, 2012; Yi et al., 2019). Further work is required to directly determine where in the brain low and high frequency speech envelopes are reflected as well as where and when the underlying neural representations give rise to the differential pattern of envelope sensitivity observed here.

## 5. Conclusion

Low and high frequency speech envelopes provide largely independent information about acoustic and phonemic features. Our results show that these envelopes are reflected in spatially and functionally distinct processes in the delta and theta band EEG. These processes are easily confounded when considering the broadband speech envelope for data analysis, but may offer independent windows on the neurobiology of speech.

## Acknowledgement

Funding: CK was supported by the European Research Council (ERC-2014-CoG; grant 618 No 646657).

